# Detecting Rhythmic Gene Expression in Single Cell Transcriptomics

**DOI:** 10.1101/2023.12.07.570691

**Authors:** Bingxian Xu, Dingbang Ma, Katherine Abruzzi, Rosemary Braun

## Abstract

An autonomous, environmentally-synchronizable circadian rhythm is a ubiquitous feature of life on Earth. In multicellular organisms, this rhythm is generated by a transcription–translation feedback loop present in nearly every cell that drives daily expression of thousands of genes in a tissue–dependent manner. Identifying the genes that are under circadian control can elucidate the mechanisms by which physiological processes are coordinated in multicellular organisms. Today, transcriptomic profiling at the single-cell level provides an unprecedented opportunity to understand the function of cell-level clocks. However, while many cycling detection algorithms have been developed to identify genes under circadian control in bulk transcriptomic data, it is not known how best to adapt these algorithms to single-cell RNAseq data. Here, we benchmark commonly used circadian detection methods on their reliability and efficiency when applied to single cell RNAseq data. Our results provide guidance on adapting existing cycling detection methods to the single-cell domain, and elucidate opportunities for more robust and efficient rhythm detection in single-cell data. We also propose a subsampling procedure combined with harmonic regression as an efficient strategy to detect circadian genes in the single–cell setting.

## 1 Introduction

The circadian rhythm is an approximate 24-hour oscillation of physiology, metabolism and behaviour that enables organisms to adapt to a changing daily environment [1, 2, 3]. At the cellular level, autonomous molecular oscillations are generated by a transcription-translation feedback loop. In *Drosophila*, for example, the CLOCK/CYCLE (CLK/CYC) heterodimer binds to the E-box to activate gene expression of period (per) and timeless (tim); the PER and TIM proteins then dimerize in the cytoplasm and translocate to the nucleus to inhibit the DNA binding activity of CLK/CYC [4]. This core clock circuit in turn drives circadian oscillations of hundreds of downstream targets. It has been reported that ∼40% of all protein coding genes may exhibit circadian oscillations in a tissue–specific manner [5], underscoring the role of the circadian rhythm in orchestrating physiological processes in multicellular organisms.

Proper temporal coordination requires synchronization of tissue–specific rhythms, which in turn requires synchronization of cell–level clocks. Loss of synchrony between organ systems and dampening of circadian rhythms has been implicated in aging and disease [6]. In older mice, for example, it has been reported that the flattened amplitude of circadian oscillation is related to deficits in long-term spatial memory due to impaired sleep [7]. Even within the same tissue, it was recently observed that the number of detectably cycling genes decreases as the animal gets older [8]. Such dampening may be due to an overall loss of amplitude, or a loss of synchrony between cells that leads to no discernible rhythm at the tissue level.

Today, single cell RNAseq profiling provides an opportunity to examine this coordination by investigating cell type–specific transcriptomic cycling. Circadian transcriptomic timeseries data (both bulk and single-cell) typically comprises samples taken every 2–4 hours over a 24–48 hour period, potentially with replicates [9]. The goal of such studies is to identify genes that are under circadian control in different conditions, providing insights into how the circadian rhythm orchestrates cellular physiology. Because these data are noisy, sparsely sampled in time, and high dimensional in the number genes, statistical tests for evidence of cycling and differential cycling remain an area of active research [10, 11, 12]. To date, however, all proposed methods were developed in the context of *bulk* RNAseq data, leaving open the question of how best to analyze circadian single-cell data. Single cell RNAseq measurements yield many more observations than bulk data (one per cell), but are much noisier due to stochastic gene expression and drop-outs; in consequence, methods designed for bulk RNAseq may not translate well to the single cell context. Below, we briefly review state-of-the-art methods for cycling detection in bulk transcriptomic data, and highlight the opportunities and challenges posed by single-cell RNAseq profiling.

Perhaps the simplest form of cycling detection is harmonic regression, in which a 24 h sinusoidal curve is fit to the data and goodness-of-fit statistics serve as an assessment of cycling. However, the noisiness of the data and the low number of replicates per timepoint in bulk transcriptomic data mean that a single outlying observation can produce a significant “cycling” component, leading to a large number of false positives when applied to bulk transcriptomic data [13]. To overcome this challenge, ARSER [14] removes linear trends in the data and smooths the data using a fourth-order Savitzky-Golay filter before the harmonic regression. In addition, there are methods that rely on Fourier analysis [15, 16], essentially conducting a discrete Fourier transform and assessing the statistical significance of the peak corresponding to a period close to 24 hours [17]. Like harmonic regression, Fourier-based methods also have inaccuracies [18, 19] due to insufficient temporal sampling (frequency and duration).

Because it has been observed that cycling genes may exhibit sharply peaked or asymmetric waveforms, non-parametric approaches have been developed to test for evidence of cycling without assuming sinusoidality [18, 19, 20, 13]. JTK-cycle [18] employs the Jonckheere-Terpstra trend test/Kendall’s *τ* rank correlation to identify smooth rise–and–fall patterns in a 24-hour window. The observed pattern of gene expression is compared to template waveforms of different phases and asymmetries (e.g. a longer rising interval than falling interval). Because corrections need to be made for the multiplicity of templates considered, JTK-cycle can lose power as the number of patterns of interest increases. To circumvent the need for predetermined waveforms, RAIN [19] was developed using the general umbrella test, a statistical test for an umbrella shape with a flexible inflection point, allowing it to accommodate to asymmetric waveforms automatically. Two alternative methods, SW1PerS [21] and TimeCycle [12], use techniques from topological data analysis to nonparametrically quantify the cyclicity of observed expression profiles. In bulk transcriptomic data, these nonparametric approaches generally outperform parametric ones. Nevertheless, it has also been shown that no cycling detection method is universally optimal, and that the best analysis method depends on both the study design (i.e. the frequency of sampling, number of periods sampled, and replicates per timepoint) and the shape of the waveforms of interest [9].

Analysis of single-cell RNAseq data presents new opportunities and challenges. The large number of cells assayed at each timepoint opens the possibility of regarding them as many replicate samples (in contrast to bulk RNAseq studies, which often have only two or three), enabling the variance between cells to inform the analysis. However, the number of cells is very large, and may vary from timepoint to timepoint. As a result, analysis methods that require a regular number of samples per timepoint (such as ARSER [14]) or those that scale poorly with the number of replicates may not be directly applicable to single–cell data. As an alternative, one can consider averaging expression levels across all cells of the same type. Methods developed for bulk transcriptomic data may then be applied to the resulting “pseudo-bulk” data. In support of this approach, it has been reported that considering cells as replicates for differential expression analysis can result in a large number of false positives due to systematic correlations between the cells coming from a common sample [22]. There, the authors found that pseudo-bulking yields more reproducible results in tests of differential expression than treating cells as independent observations. However, it is not known to what degree pseudobulking may enhance the specificity of cycling detection (by reducing false positives) vs. reducing its sensitivity (by lowering the number of replicates), and there remains no guidance on the best way to analyze single-cell circadian timeseries data.

The goal of this work is to evaluate the performance of various existing approaches applied to a circadian timecourse single-cell RNAseq dataset of *Drosophila* brain tissue [23], as well as to synthetic data. We applied a number of different methods (including JTK-cycle and RAIN) as well as a variety of application approaches (i.e., treating cells as distinct samples or averaging to create a pseudo-bulk) and assessed the computational efficiency, reproducibility, and robustness to noise of the algorithms. Our results suggest that methods designed for circadian detection in bulk RNAseq data may not be optimal for single-cell data, and suggest opportunities for new approaches.

## 2 Materials and Methods

### 2.1 Data acquisition

We use publicly available single–cell RNAseq data from Ma *et al.* [23](https://www.ncbi.nlm.nih.gov/geo/query/acc.cgi?acc=GSE157504) in our study. The data comprise four circadian time-series of fly brains, with two time-series collected under each of two conditions (12h light/dark [LD] and dark/dark [DD]). Each series comprises six samples collected every four hours. In total, 4671 cells were assayed (Supplementary Table 1). Normalized data was obtained as a Seurat object [24, 25] containing all relevant metadata. The original authors annotated these into 39 clusters, of which 17 were assigned to known cell types (Supplementary Table 1). In the analyses that follow, we used the normalized counts of the LD cells as input to the cycling detection algorithms. Details of the preprocessing may be found in the Supplement.

### 2.2 Creation of pseudo-bulk expression profiles

In addition to considering single cells as individual observations, we also constructed pseudo-bulk expression profiles for each cell type (cluster). Pseudo-bulk profiles were constructed from the normalized expression matrix *X* ∈ **R***^g×c^* where *g* denotes the number of genes and *c* the number of cells for the cell-type of interest. Each column in *X* corresponds to a cell with a time stamp *t_i_* ∈ {*t*}, where {*t*} is the set of sampling times. The pseudo-bulk expression profile for gene *g* at time *t* is the average across the cells in a given cell type with timestamp *t_i_* = *t*,

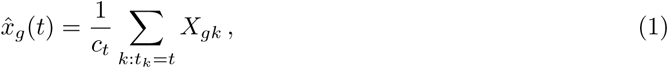

where *c_t_* denotes the number of cells sampled at time *t*. Pseudo-bulk temporal profiles were calculated per Eq. 1 for the four largest clusters (Figure 4).

### 2.3 Application of cycling detection algorithms

JTK-cycle [18], RAIN [19], and harmonic regression were applied to single cell data using the recommended parameter settings (see Supplement), treating each cell as an independent sample. Additionally, we applied JTK-cycle, RAIN, harmonic regression, and ARSER [14] to the pseudo-bulk data. ARSER, harmonic regression, RAIN and JTK-cycle were all implemented using their respective R packages [18, 26, 19, 27]. We excluded from our analysis SW1PerS [21], which does not return a *p*-value, and TimeCycle [12], which requires at least 18 time points to construct the cycle.

### 2.4 Systematic data contamination

In addition to testing the performance of the algorithms against the original data, we wanted to explore whether a systematic artefact affecting a single time-point would generate false–positive results. In [22], it was demonstrated that differential expression analysis is prone to false positives when cells are considered replicates due to the fact that a small artefact would be amplified by the large number of cells. Would the same hold true for cycling detection?

To examine this possibility, we identified genes that showed no evidence of cycling, and systematically contaminated the data by elevating the expression level of genes at specific timepoints to explore whether the algorithms would erroneously identify it as cycling. We restricted this analysis to the largest cell cluster (1:DN1p_CNMa) collected under the LD condition. At a given timepoint, we increased the expression level for each gene in each cell by an amount proportional to the gene’s average expression across all timepoints. For each round of analysis, we increased the expression level of all cells for a single time point, mimicking an batch effect affecting a single scRNA-seq collection. That is, for each gene *X*(*c, t*) measured in cell *c* at timepoint *t*, we increase its expression to

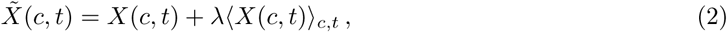

where 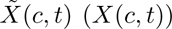 denotes the contaminated (original) normalized count for gene *X* in cell *c* at time *t*, ⟨·⟩*_c,t_* denotes an average over all cells (from the same cell type) at all timepoints, and *λ* is a parameter that controls the extent of the contamination. Non-cycling genes selected for synthetic contamination were chosen as those that were detectably expressed in at least half of the cells and had a harmonic regression *p >* 0.9 (treating cells as replicates). This yielded 59 “non-cycling” genes.

### 2.5 Generating synthetic data

In addition to analyzing real single–cell data, we generated synthetic data following the steps outlined previously [28]. Briefly, we modeled counts sampled at each sampled time point (t) to follow a negative binomial distribution:

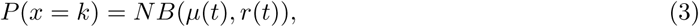

where *µ* and *r* are the mean and dispersion respectively. The variance of a negative binomial distribution is 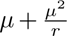, which increases when *µ* increases or when *r* decreases. Cycling signals are generated by varying *µ* with time. In order to ensure that our synthetic data would be biologically representative, we selected the 20 genes with the lowest harmonic regression *p*-values from the real data and 20 genes from the aforementioned “non-cycling” genes to use as templates for varying *µ*. The expression level of each gene was first normalized such that it ranges between 0 to 100. For each synthetic signal (Figure S5), *µ*(*t*) is chosen to be the mean expression of a selected template gene at the given time point, plus a time independent expression offset (*µ*_0_). Namely, at a given time point *t*, *µ*(*t*) for a given gene is set to be:

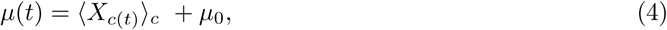

where ⟨*X_c_*_(_*_t_*_)_⟩*_c_* denotes the mean expression level of the template at time *t*. The addition of the time-independent offset (*µ*_0_) allows us to examine if methods can detect oscillations whose amplitudes are small relative to the mean (and hence also the variance).

With this, we generated synthetic single cell RNA-seq data by sampling from negative binomial distributions for 12 time points, with 5 samples from time point. This choice accommodates all tested methods, as some cannot handle large datasets, and others cannot handle uneven replicates. In total, we selected a total of five and seven levels of *µ*_0_ and dispersion, ranging from 0.1 to 500 and 0.001 to 5 respectively, generating an ensemble 1400 synthetic gene expression time series. In the fly scRNA-seq data, the estimated dispersions [24] range from 10*^−^*^6^ to 8, with a mean and median of 0.43 and 0.2 respecively. Our simulated dispersions span beyond the 5^th^ (0.014) and 95^th^ (1.72) percentile of those observed in the real scRNA-seq data.

## 3 Results

### 3.1 A framework for evaluating cycling detection reproducibility in singlecell RNAseq datasets

The large number of observations (cells) obtained at each timepoint in a single-cell RNAseq experiment permits a novel approach for evaluating the reliability of cycling detection methods. The approach is based on the following conjecture: if a gene is truly under circadian control in a certain cell type, it should be detected as cycling reliably even when considering only a subset of the cells at each timepoint. We note that for most cell-type clusters, even taking half of the observed cells yields more replicates than are typically found in bulk circadian transcriptomics studies (where there are commonly ≤3 observations per timepoint). We thus compute the reliability of a cycling detection method as follows. For each cell-type at each timepoint, we select at random half of the cells as “sub-experiment” 1, and consider the rest as “sub-experiment” 2. We then apply the cycling detection algorithm to both sub-experiments, yielding for each sub-experiment the genes detected as cycling. Ideally, the intersection of those sets should be complete, with the same genes detected as cycling (or not) in each sub-experiment. We use the size of the intersection *N_∩_* over the number of cyclers detected when all cells are used *N*_full_ as a measure of reliability. The choice of developing this measure rather than using existing measures such as the Jaccard index (*N_∩_/N_∪_*, where *N_∪_* is the union of the cyclers in the two sub-experiments) is that the union of the cyclers detected in the two sub-experiments *N_∪_* are only a small fraction of the cyclers detected when all cells are used (*N_∪_*≪*N*_full_). Since typical analysis of single–cell data would use all cells, we took *N*_full_ to be the basis for comparison. The same procedure can also be used with pseudo-bulk data; in this case, division into the two sub-experiments is done before Eq.1 is calculated.

In our tests, we performed 10 random splits of the data into the two sub-experiments to gather statistics for *N_∩_*. Higher *N_∩_/N*_full_ indicates greater reproducibility of the cycling detection, even when the data are subsampled.

### 3.2 Computational efficiency

We first tested how run-time scales with the total sample size, ranging from ∼10 to 200 cells, when treating each cell as an independent observation (i.e. without pseudo-bulking). We focused on three methods—JTK-cycle [18], RAIN [19], and harmonic regression [27]—chosen because of their popularity and their ability to handle uneven replicates. The latter is a crucial feature, since the number of cells is likely to vary per time-point in single-cell data; as a result, we exclude methods (such as ARSER) that require a consistent number of observations at each timepoint. We then considered the computational cost of running these algorithms without pseudo-bulking.

In contrast to what was previously reported [13], we observed that RAIN can actually be faster than JTK-cycle when the sample size (i.e. number of cells) is small. However, as sample size increases, RAIN becomes slower than JTK-cycle as expected. These differences in efficiency are likely due to implementation details, rather than the complexity of the underlying test; we derived the theoretical scaling relationship as having O(*m*^2^) computational complexity in the number of samples *m* in both cases (see Supplementary Information). However, we observe empirically that the measured run time for both JTK-cycle and RAIN deviates considerably from the estimated quadratic complexity (Figure 1). Our empirical benchmarking suggests that for a moderate single cell dataset with only 1000 cells, JTK-cycle (RAIN) will take at least seven hours (sixteen days) to complete on a MacBook Pro with 2.9GHz dual-core Intel Core i5 and 8G memory. In particular, both implementations appear to scale exponentially, which may make them unfeasible for application to single-cell data without pseudo-bulking. The discrepancy between the theoretical computational complexity and the empirical performance also suggests that these algorithms’ efficiency may still be improved. Harmonic regression, as expected, scales linearly in the number of observations, and takes *<* 10s to compute even for very large clusters.

**Figure 1:**
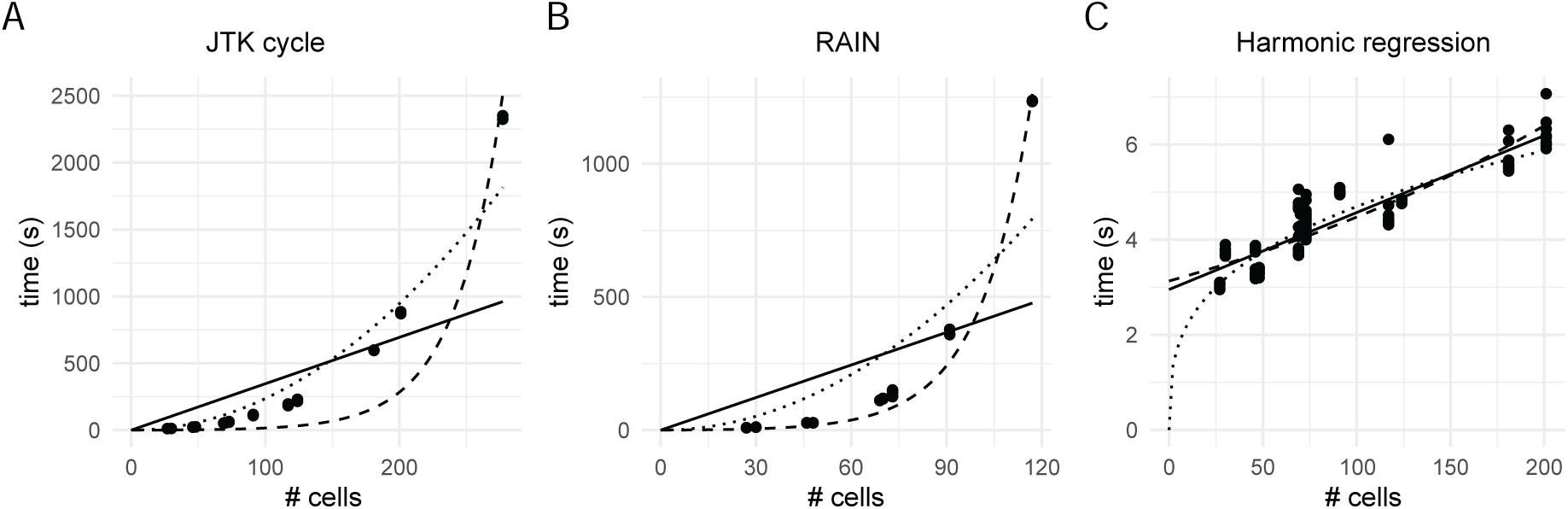
Scaling relationship between the number of samples and run time for (A) JTK-cycle, (B) RAIN, and (C) harmonic regression. Trend-lines were fitted for linear (solid), quadratic (dotted) and exponential (dashed) computational complexities. Note the difference in *y*-axis scales.

### 3.3 Reliability of cycling detection in single-cell data

We next explored the reliability of cycling detection in sc-RNAseq data using the concordance framework described above. As in the previous section, we applied JTK-cycle, RAIN and harmonic regression to eight cell types within the Ma data that contained at least two cells at each time point, and which were sufficiently small to analyze given the efficiency attributes noted above. Our choice of methods is motivated both by their popularity [29, 8, 23, 30] and their ability to handle replicates and run efficiently when sample size is low [13]. For all methods tested, we consider a gene to be cycling if the *p*-value is less than 0.05. We compared how these three methods differed in the number of cycling genes detected and in their subsample concordance. Figure 2 shows the number of genes detected as cycling when no sub-sampling is done (*N*_full_, panel A); the number of genes in the intersection of two sub-experiments when subsampling is done (*N_∩_*, panel B); and their ratio (panel C).

**Figure 2:**
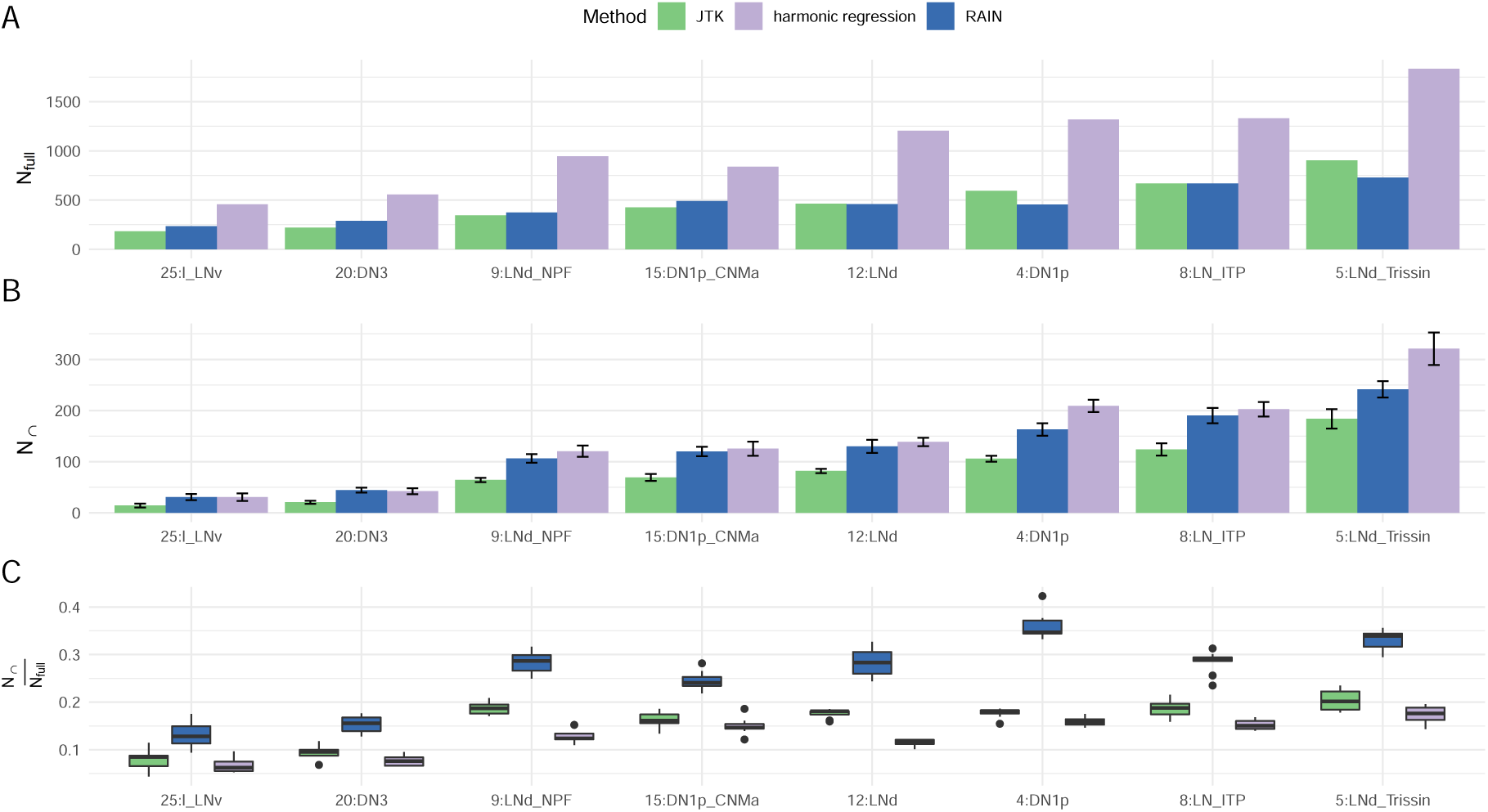
A: The number of cycling genes detected using all cells, *N*_full_. B: The average number of cyclers detected in both subsamples across 10 trials, *N_∩_*. C: The ratio of *N_∩_* to *N_full_*, the proportion of consistently–detected sub-sample cyclers relative to those found using all cells. Tests were conducted in the largest eight labeled cell clusters that RAIN can handle. JTK-cycle and harmonic regression were run in parallel. Error bars indicate standard deviation across the 10 subsamplings. Cell types are labeled as clusterID:cell type identity, obtained from the dataset authors’ original labeling [23]. For example, 25:l_LNv refers to cluster 25, large ventral lateral neurons. 20:DN3: dorsal neuron. 9:LNd_NPF: NPF expressing dorsal lateral neurons. 15:DN1p_CNMa: CNMa expressing posterior dorsal neuron. 12:LNd: dorsal lateral neurons. 8:LN_ITP: ITP expressing lateral neurons. 5:LNd_Trissin: Trissin expressing dorsal lateral neurons.

Interestingly, while it has been reported that RAIN is less conservative than JTK-cycle in bulk data [9, 19], we observe that the number of detected cycling genes by these two methods tend to be similar when cells were considered as replicates (Figure 2). We then conducted subsampling (as described above) to examine the reproducibility of genes detected as cycling in both subsamples. Interestingly, though harmonic regression considers more genes to be cycling compared to JTK-cycle and RAIN, all the obtained *N_∩_* tended to be similar (Figure 2). Lastly, we looked at the ratio between *N_∩_* and *N*_full_ and observed that RAIN had the highest subsample concordance in all tested cases, around two times greater than that from both JTK-cycle and harmonic regression.

We note that the above analysis used unadjusted *p*-values. To examine if adjusting *p*-values for multiple hypotheses increases subsample concordance, we repeated above analysis using FDR values computed with the Benjamini-Hochberg method, setting *FDR <* 0.05 as the cycling detection threshold. Interestingly, we observed a general *decrease* of subsample concordance (Figure S1) with this more stringent criterion, suggesting that using the FDR is insufficient to robustly identify circadian genes.

We then examined whether the various methods identified the same genes as cycling both across subsamples and across methods (Figure 3 and Supplemental figures S2, S3). Harmonic regression yielded many subsample-specific cyclers (i.e. genes detected as cycling in one but not the other of N1, N2, shown as the first two sets in the upset plot), suggesting that the harmonic regression *p*-value alone may generate a large amount of false–positives. However, we observe that considering the *overlap* in the sub-experiments mitigates this issue. For the genes detected by harmonic regression in *both* sub-experiments N1 and N2 (ie, sets with overlap in HR-N1, HR-N2 in Figure 3), the majority are also identified by RAIN and JTK-cycle as well. Additionally, we again observed that adjusting *p*-values does not increase the agreement between each experiment (Figure 3).

**Figure 3:**
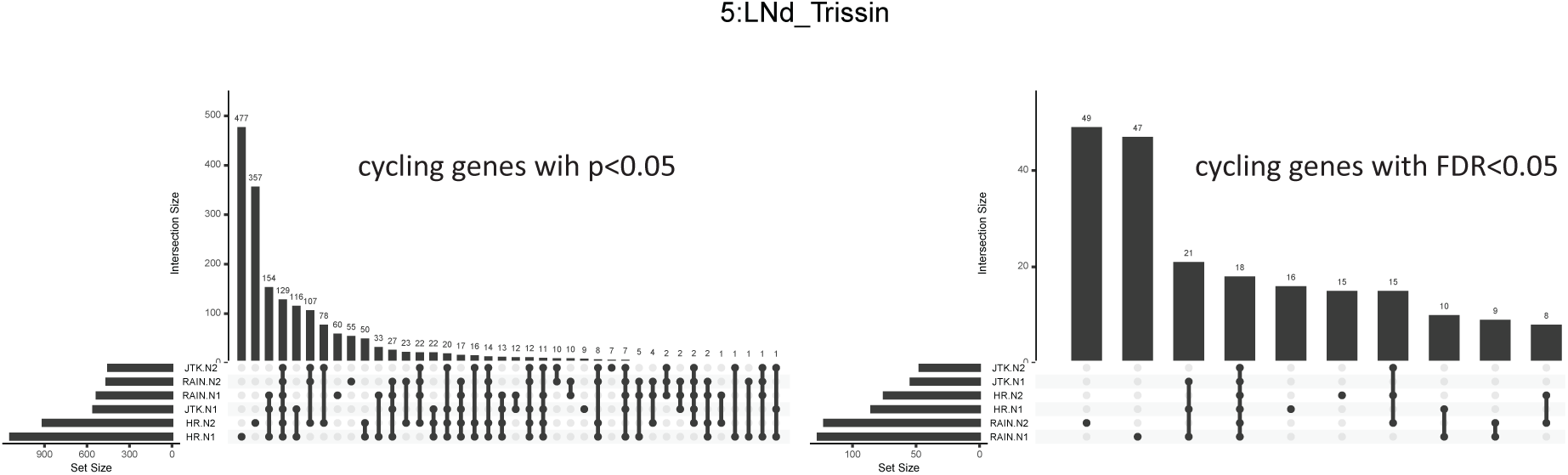
UpSet plots generated from Trissin expressing dorsal lateral neurons (5:LNd_Trissin) showing the intersection of detected cycling genes, with either *p*-values or FDR–adjusted *p*-values less than 0.05, by harmonic regression, RAIN and JTK-cycle in one instance of the two sub-samples. Here, HR indicates harmonic regression; JTK = JTK-cycle; N1 = sub-experiment 1; N2 = sub-experiment 2.

Importantly, this suggests that looking for genes *consistently* detected as cycling across two subsamples using a fast algorithm (harmonic regression) may be a reliable and efficient alternative to more computationally intensive methods. That is, while harmonic regression alone identifies many genes as cycling (some proportion of which may be false positives), it is unlikely that a gene will be spuriously identified as cycling in two subsamples. As shown in Figure 3, that are called as cycling in two subsamples of the data tend also to be ones that would be called as cycling using other, less efficient methods.

### 3.4 Cycling detection using pseudo-bulk data

The problems of large run-times and uneven replicates can also be circumvented by first averaging across cells of a given cell type within each timepoint, yielding a single “pseudo-bulk” gene expression value for each cell type at each timepoint. Cycling detection may then be applied to the resulting pseudo-bulk data, as was done in a dataset of mouse SCN neurons [30]. To examine how pseudobulking affects cycling detection, we applied JTK-cycle and harmonic regression to pseudo-bulk data and compared it against the results obtained using cells as replicates. As an additional comparison, we also conducted cycling detection on pseudo-bulk data using ARSER (which cannot handle uneven replicates) and RAIN (which is computationally inefficient in the non-pseudo-bulk setting, Figure 1B). We observed that the number of cycling genes (*N_full_*) detected using pseudo-bulk data was similar for JTK-cycle and ARSER. RAIN identified the largest number of rhythmic transcripts, approximately twice that of JTK-cycle and ARSER, and ected from previous studies that also noted RAIN’s permissiveness [13]. For JTK-cycle and harmonic regression, where we can directly compare the outcome of treating cells as replicates or as a pseudo-bulk tissue, we observed that both methods detected more cycling genes when cells were treated as replicates, as expected given that smaller *p*-values obtain from greater sample sizes (Figure 4A). Tests in synthetic data, described below, provide further insight into this phenomenon.

**Figure 4:**
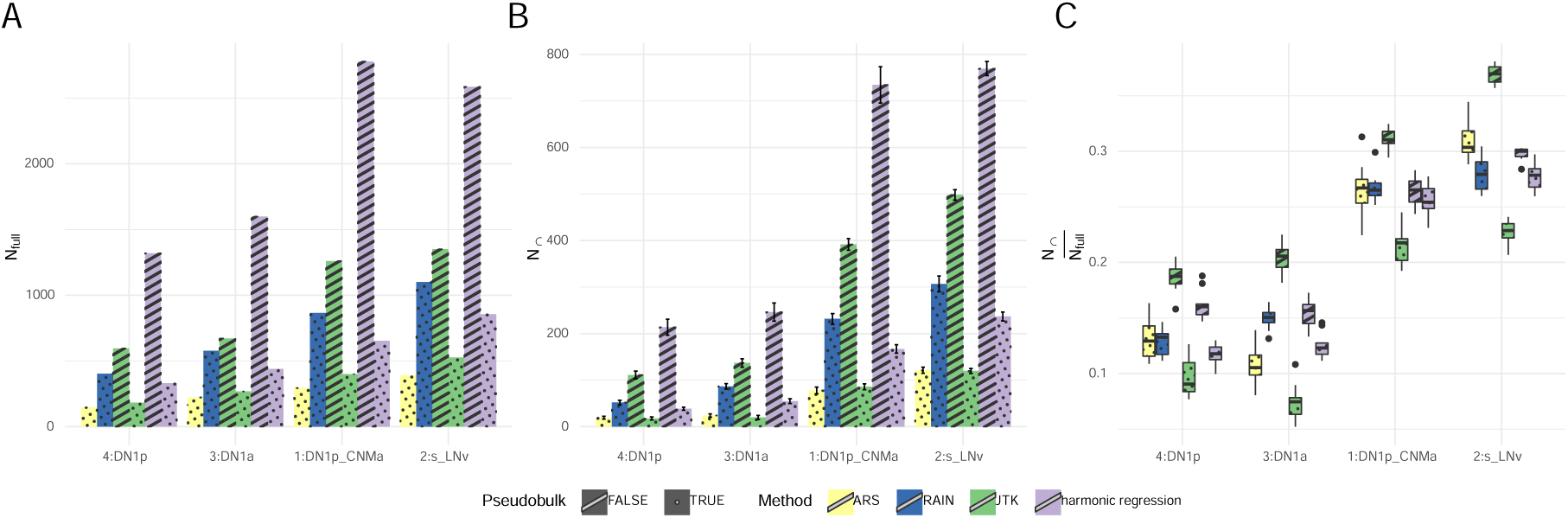
A: the number of cycling genes detected using each method. B: the number of genes detected as cycling in both samples in each method. C: subsample concordance of each method. DN1p: posterior dorsal neurons. DN1a: anterior dorsal neurons. s_LNv: small ventral lateral neurons.

In addition, we split our data into two subsamples as described previously, both containing cells collected at all of the time points as required by JTK-cycle and RAIN. With this, we conducted both the pseudo-bulking and the subsequent cycling detection using all samples (“full” data), as well as the first and second subsample separately. We then counted the number of genes in the intersection of those considered cycling in both subsamples (*N_∩_*), and quantified *N_∩_/N_full_* as a measure of the concordance. From our analysis, we observed the lowest subsample concordance when JTK-cycle was applied to the pseudo-bulk (Figure 4B,C).

### 3.5 Impact of systematic contamination

While we have shown that considering cells as replicates can reliably detect more cyclers than pseudobulking, it has been reported that methods designed to conduct single cell differential expression analysis where cells were considered as replicates tend to overestimate the number of differentially expressed genes, due to the fact that the cells from a given sample are not truly independent observations [22]. To test whether the same idea holds true for circadian detection algorithms, we first synthetically increased the expression of non-cycling genes at individual timepoints (see Materials and Methods) and applied harmonic regression on data with and without pseudo-bulking. In each round of our analysis, we elevated the expression of all cells collected at the selected time point. As expected, the estimated phase of these non-cyclic gene converged to the time of elevated expression (Figure 5A). Additionally, we observed that *p*-value decreases as the magnitude of gene expression elevation increases; consequently, a gene will have a higher chance of being falsely called as cycling when this type of systematic noise is high. Though *p*-value decreases with or without pseudo-bulking, we observed zero false–positive cyclers when pseudo-bulk data is used, suggesting that pseudo-bulking is robust to technical artefacts such as elevated expression at a single time point (Figure 5A).

**Figure 5:**
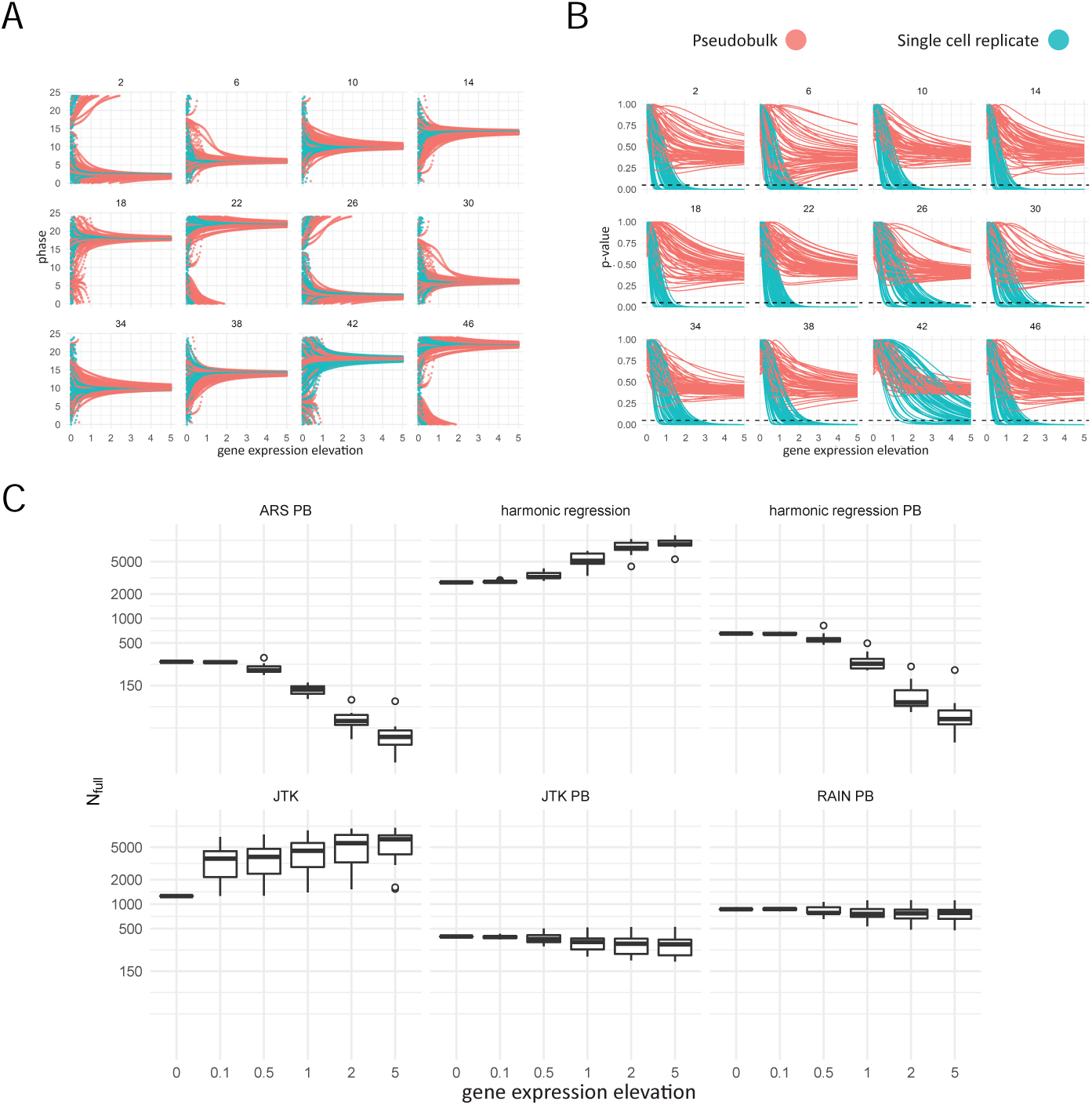
A: Harmonic regression estimated phase of each non-cyclic gene as a function of gene expression elevation. The number of each subplot indicates the timepoint which was subject to gene expression elevation. B: Harmonic regression *p*-value of each gene as a function of gene expression elevation. The dashed line indicates *p* = 0.05. C: The number of cyclers detected at each level of gene expression elevation.

We next examined the effect of elevating the expression of all genes (including those considered cycling) for all methods, using the DN1p_CNMa cells as an example. In general, we observed that the number of detected cyclers increases as a function of the magnitude of contamination for methods that consider each cell as a single replicate (JTK-cycle and harmonic regression) (Figure 5C). As expected, the number of detected cyclers did not increase indefinitely for JTK-cycle as it only depends on the rank order of gene expression, which eventually stops changing and does not impact the outcome of the cycling detection algorithm any further. For methods where pseudo-bulk data was used as input, we observed a decreasing number of detected cyclers as a function of contamination, where ARSER and harmonic regression experienced the greatest amount of loss of detected cyclers (Figure 5 A). Closer inspection reveals connections between the methods. When looking at how the number of total detected cycling genes changed as a function of noise magnitude and the affected time-point, we observed that JTK-cycle and RAIN, both using the Jonckheere-Terpstra test, behaved similarly when applied to pseudo-bulked data (Figure S4). Similar observations were also made when harmonic regression and ARSER were applied to pseudo-bulked data (Figure S4) and when harmonic regression and JTK-cycle were applied considering each cell as a replicate (Figure S4).

In total, we observe that considering cells as replicates can give rise to false positive cyclers in the presence of strong experimental artifacts. In contrast, pseudo-bulking affords protection from this effect, but at the cost of detecting fewer cycling genes.

### 3.6 Cycling detection on synthetic data

Using real data, we have shown that low subsample concordance is a common problem for all cycling detection methods both when cells are treated as replicates and when pseudo-bulk profiles are constructed. However, the construction of pseudo-bulk data offers protection against outlier datapoints, as may arise from batch effects. Additionally, however, single cell data are known to be noisy. To explore the sensitivity of each method to gene expression noise, we constructed synthetic data for cycling and non-cycling gene expression profiles (Figure S5). By modeling count data using negative binomial distributions, we tested how each method performed under different levels of gene expression noise (variance), controlled jointly by the dispersion and mean expression (see Materials and Methods for detail). Consistent with previous results, we observed that the *p*-values computed from non-parametric methods (JTK-cycle and RAIN) are larger than that those from parametric methods (harmonic regression and ARSER).

At low variance (high dispersion), we find that *p*-values from pseudo-bulk data tend to be higher than that those when cells were considered as replicates (Figure S6, left columns), leading to fewer detected true cyclers (Figure S7A) but also fewer false positives (Figure S7B). This is consistent with previous studies indicating that *p*-values for differential expression are artificially low when considering cells as replicates, and suggests that pseudobulking offers protection from false positives.

When variance is moderately high, however, we observed the opposite effect, where in pseudobulk produced *lower p*-values than the independent cell analysis (middle columns in Figure S6), leading to a small “bump” of false positives in Figure S7B. For example, we observed that RAIN with pseudobulk can produce false positives when *µ*_0_ = 1 and dispersion = 0.05 (Figure S6B, S7B). Upon closer inspection, we observed that this phenomenon originates from genes whose expression is largely zero. In this case, pseudobulking introduces false positives by generating outlier-driven oscillations in the mean. We have observed genes with similar expression pattern in real data as well (Figure S8). Once the variance is sufficiently high (rightmost columns in Figure S6, S7) no method reliably detects cycling genes.

### 3.7 Comparison to bulk RNA-seq data

We next examined whether genes detected as cycling in bulk RNAseq samples of fly neurons are also picked up as cycling in the single–cell data. We assembled a list of published circadian genes in the fly clock neurons [31], originally identified using a combination of JTK-cycle and Fourier analysis, which contains three distinct clock neuron types, LNv (ventral lateral neurons), LNd (dorsal lateral neurons), and DN1 (dorsal neurons). We examined the scRNA-seq *p*-values for the genes on/off those lists for the clusters of cells that corresponded to the neuron types assayed in the bulk data (Figure S9). In Figure 6, we plot ROC curves for “correctly” classifying a gene as a function of the single–cell *p*-value, where here “correctness” is defined by whether the gene was reported as cycling in the bulk. (Note, however, that the bulk data may also have spurious or missed cyclers; discrepancies with the single-cell data may be due to *correct* calls by the single-cell analysis.) We observed that RAIN has the best performance at discriminating bulk cyclers and non-cyclers, with or without pseudo-bulking (Figure 6). Interestingly, JTK-cycle, which uses the same underlying statistical test as RAIN, has the worst overall performance. The origin of this difference is not clear.

**Figure 6:**
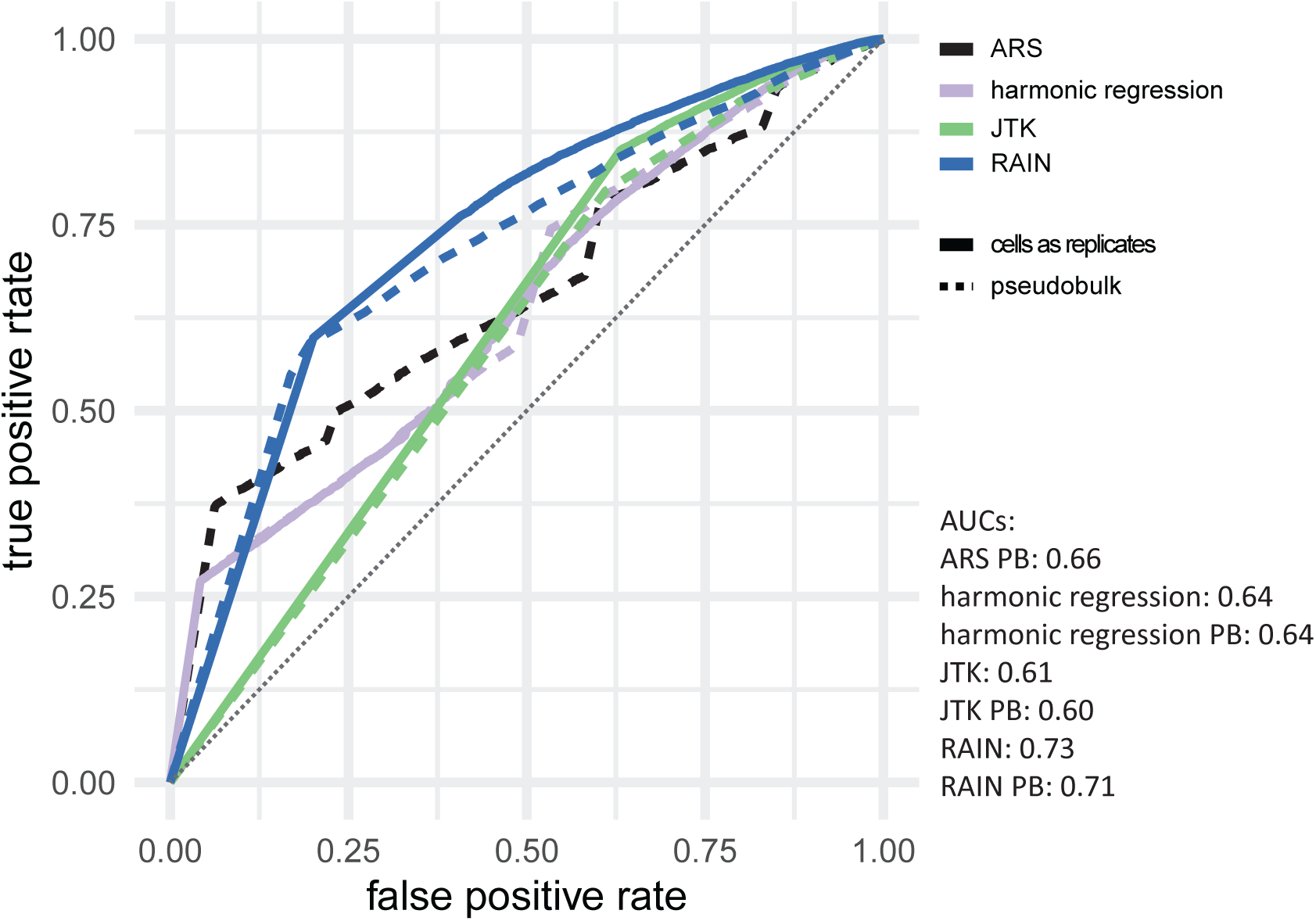
ROC curves for identifying reported bulk circadian genes using single cell RNA-seq data, as a function of *p*-values from various cycling detection algorithms. The grey dotted line indicates *y = x*.

## 4 Discussion

We evaluated the performance of four cycling detection algorithms on single cell transcriptomic timeseries data, with and without pseudo-bulking. Our analysis indicates that existing cycling detection algorithms scale poorly with replicate size, and showed low subsample concordance in general.

While the problem of long run times can be circumvented by pseudo-bulking, we find that cycling detection on pseudo-bulked data identifies fewer cycling genes in comparison to treating each cell as a replicate. To test whether this difference in the number of cyclers is meaningful, we conducted subsample concordance analysis by splitting the data into two subsamples to see how the number of cycling genes detected in the full data compares to that of the intersection of the two subsamples. By conducting this analysis on four major clock neuron clusters, we observed that pseudo-bulking significantly reduced 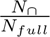, with greater *N_∩_* when using cells-as-replicates. This suggests that the cyclers detected after pseudo-bulking may be less reproducible (more differences between sub-experiments) than those detected when treating cells as replicates.

Our observations with simulated data agree with those from real data, where in general, nonparametric methods generate higher (less significant) *p*-values than parametric methods. Here, we find that pseudobulking has an interesting effect. At relatively low levels of noise (dispersions ≥ 1 [leftmost columns] in Fig S6), pseudo-bulking generates higher (less significant) *p*-values than considering cells as replicates. This offers protection against false positives, exemplified by Fig S7B and our previous results where pseudobulking offers robustness to artificial gene expression elevation. However, it comes at the cost of a higher false negative rate, exemplified by Fig S7A as well as our comparison to bulk RNA-seq in which the pseudobulk single-cell analysis was slightly less able to identify bulk cyclers than the single-cell analysis considering cells as replicates was (slightly lower AUCs in Figure 6). At moderate noise levels, however, the effect is the opposite, with lower *p*-values obtaining from pseudobulk (middle columns in Fig S6) and an increase in false positives (“bumps” in the center of Fig S7B). Further investigations revealed that this is due to spurious appearance of cycling due to outliers in genes that in general are not expressed (see examples in Fig S8). Altogether, this analysis suggests that pseudobulking should be undertaken with care in the context of cycling detection.

Our analysis also points to a possible method to address the problem of long run times while treating cell as replicates to maintain high 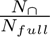 : specifically, using the fast (but suboptimal) harmonic regression on two randomly selected subsets of cells, and then considering the overlap of the results. Considering the overlap of the subsets mitigates the false–positive issues that affect harmonic regression. In Fig 3, we demonstrate that using this overlap yields the same genes as would be detected via other methods.

However, care should still be taken when treating cells as replicates. In single cell differential expression analysis, it has been shown that treating cells as replicate can lead to false positives. To investigate whether the same happens in cycling detection algorithms, we artificially contaminated our data by elevating the expression at a single time point, mimicking what might happen if a single sample in a circadian time–series was affected by a systematic artefact. We then compared how pseudobulking impacts the outcome of commonly used circadian detection algorithms. When all genes (both cycling and non-cycling in the original data) were contaminated by a systematic offset of one time– point, we found that the number of detected cyclers increased with the amount of contamination when cells were considered as replicates, suggesting that treating cells as replicates may result in false positives due to experimental artefacts. On the other hand, the number of cycling genes decreased with the amount of contamination when pseudo-bulk data was used, suggesting that pseudo-bulking may provide protection from false detection of cyclers, albeit at the expense of detecting fewer true cyclers. Together, these observations suggest that choices should be made based on the analytical goals of the study. When one wants to investigate collective patterns, such as phase distributions of cycling genes, it may be beneficial to consider each cell as replicates despite the fact that it may include false positives. On the other hand, if the need is to identify the most robust cycling genes, it is better to conduct cycling detection on pseudo-bulked data.

We acknowledge that a limitation of the present study is that we confined our analysis to preprocessed data. Different normalization and clustering methods will inevitably alter cell type assignment and the normalized expression level for each gene, affecting downstream analyses. While optimizing the normalization for the purpose of cycling detection was beyond the scope of our benchmarking, it would useful to investigate this in future work.

Finally, our analysis highlights the importance of computing subsample concordance. By looking at the agreement between detected cyclers from two subsamples, we noted poor subsample concordance from all tested methods. This provides an opportunity for designing and evaluating new methods for cycling detection designed specifically for single cell datasets.

## 5 Acknowledgements

This work was supported by NSF grant DMS-1764421, Simons Foundation grant 597491, and NIH grant R01AG068579.

## 6 Code and Data Availability

Code for our analysis is available on bitbucket (https://bitbucket.org/biocomplexity/singlecell_ benchmark). Annotations for the circadian single-cell RNA-seq (GSE157504, https://www.ncbi.nlm. nih.gov/geo/query/acc.cgi?acc=GSE157504) [23] may be found at https://github.com/rosbashlab/ scRNA_seq_clock_neurons.

## 7 Author Contribution

XB and RB designed the research. DM and KA generated the data. XB generated synthetic data, developed code for all benchmarking analyses, and analyzed the data. XB and RB wrote the paper.

## 8 Supplemental material

### 7.1 Data preprocessing and clustering

Preprocessed data and cell type information was generated as described by Ma et al [23]. Briefly, single cell data from six time points (two experimental replicates, hence twelve in total) were integrated the data on a per time point basis using Seurat. Anchor genes were identified using the FindVariableFeature function, and only those considered to be features at all time points were selected. Prior to integration, the data was first processed using the scTransform function. Raw counts were normalized using the NormalizeData function in Seurat with default parameters. Integrated data were then reduced to 2D using t-SNE, from which cluster identities were assigned. Full preprocessing details may be found in [23], with code and cell-type annotations available from https://github.com/rosbashlab/scRNA_ seq_clock_neurons. For all of our analysis, we used normalized counts generated by Ma *et al*. to ensure correspondence with results reported in [23]. A copy of the resulting normalized data can be found at https://bitbucket.org/biocomplexity/singlecell_benchmark.

### 7.2 Cell type selection for figures 2, 4

Not all cell clusters were used for Figures 2–4. Our choices are described here.

For Figure 2, we compared how different cycling detection methods perform when considering each cell as a replicate. To ensure the data is compatible with RAIN, JTK-cycle and our subsampling scheme, we combined both replicates so that each time point has at least two cells. To ensure RAIN can be applied with a tractable amount of time, we excluded clusters with more than 120 cells. The selected clusters have sample size ranging from 30 to 117, covering the full range of sample size in this dataset.

For Figure 4, we compared how different cycling methods perform when considering each cell as a replicate and when pseudo-bulk expression profile is generated. We concatenated the two replicates so that the time series spans two periods (corresponding to the actual collection of the data) and selected all cell types with at least two cells at each time point for subsampling.

### 7.3 Cycling detection

Because the two experimental replicates from Ma et al. were collected consecutively, the data were concatenated to a time series with twelve time points. Consequently, the pseudo-bulk time series also consist of twelve data points, and similarly for the subsampled data (constructed such that each subexperiment has at least one sample from each one of the twelve time points).

We applied four cycling detection methods:

- JTK-cycle is implemented using *JTK*_*CY CLEv*3.1*.R*. The period was set to 6 and the sampling interval was set to 4 to match the experimental procedure. When considering single cells as replicates, we also supplied the number of samples for each time point.
- Harmonic regression is implemented using the harmonicRegression package in R [27] inputting only time and the expression matrix.
- RAIN is implemented in the R package [19]. It takes as input the expression matrix ordered by sampling time and the sampling interval *deltat*, set to 4 to match the experimental design.
- ARSER is implemented using the metacycle package in R [26], whose only required input is the expression matrix and sampling time.

For the purpose of our analysis, a gene is considered to be cycling if its raw *p*-value is less than 0.05. We also repeat the analysis using FDR–adjusted *p*-values.

### 7.4 Derivation of theoretical computational complexity

#### 7.4.1 Mann-Whitney U Statistics

Both JTK cycle and RAIN are non-parametric methods that are built upon the Mann-Whitney *U* statistics. Let (*X*_11_*, …, X*_1_*_m_*),… (*X_T_* _1_*, …, X_T m_*) be a set of *T* time–associated observations following probability distributions *P*_1_(*x*), …, *P_T_* (*x*) for timepoints 1–*T* . For the purpose of deriving computational complexity, we have assumed that each time point contains the same number of cells *m* without loss of generality. The Mann-Whitney test tests the null hypothesis that *P_i_*= *P_j_*(*i_i_*; *j*) by computing the *U* statistic between sample *i* and sample *j* as

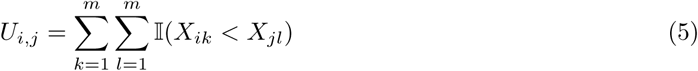

It is easy enough to see that *U_i,j_* should have a computational complexity of O(*m*^2^) and therefore the computational complexity of **U** = (*U*_1_,_2_*, …, U_T−_*_1_*_,T_*) should be approximately O(*T* ^2^ × *m*^2^). Since in real datasets *T* ≪ *m* and the number of time points do not change from one cluster to another, our expected observed computational complexity, when changing sample size, should also be O(*m*^2^).

#### 7.4.2 Jonckheere-Terpstra Test

JTK cycle employs the Jonckheere-Terpstra test, which is an extension of the Mann-Whitney U statistics to the case of having more than two samples. It tests for the presence of monotonic trend with the test statistics *s*, defined as

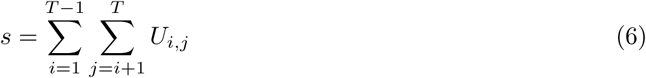

Again, it is clear that the computational complexity for *s* should be O(*m*^2^ + *T* ^2^). When the number of time points is fixed, the computational complexity scales quadratically with sample size.

In JTK cycle, the rising and falling part of the oscillation is tested for each a pre-determined waveform (e.g., sine waves of differing phases), resulting in a complexity of O(*km*^2^), where *k* is the number of templates. As *k* is fixed and *k* ≪ *m*, the complexity will scale approximately O(*m*^2^).

#### 7.4.3 General Umbrella

To overcome the JTK-cycle’s loss of power when considering multiple waveforms, RAIN uses a variation of the general umbrella which tests for the presence of an umbrella shape, which can be formally written as

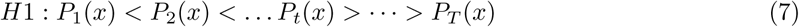

Here, *t* is a predetermined inflection point. The test statistics of the general umbrella is the sum of two Jonckheere-Terpstra statistics

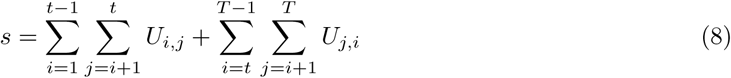

RAIN tests for all potential inflection points *T*, yielding a computational complexity O(*Tm*^2^). Since *T* is fixed by the experimental design, the complexity scales with the squared number of cells.

---

**Figure S1:**
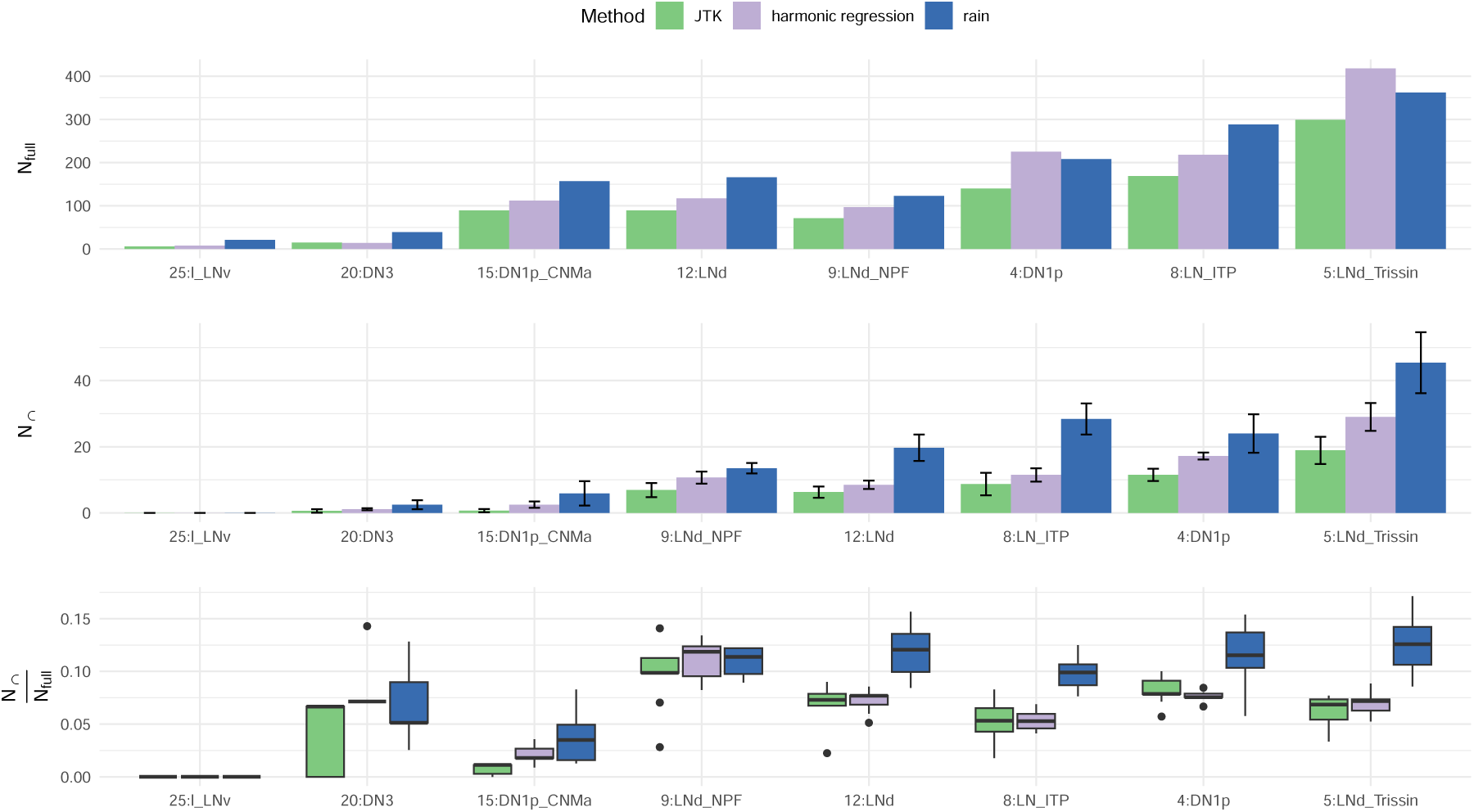
Top: The number of cycling genes detected using all cells, *N*_full_. Middle: The average number of cyclers detected in both subsamples across 10trials, *N_∩_*. Bottom: The ratio of *N_∩_* to *N_full_*, the proportion of consistently–detected sub-sample cyclers relative to those found using all cells. Error bars indicate standard deviation across the 10 subsamplings. A gene is considered cycling if its BH corrected *p*-value (i.e., FDR) is less than 0.05.

**Figure S2:**
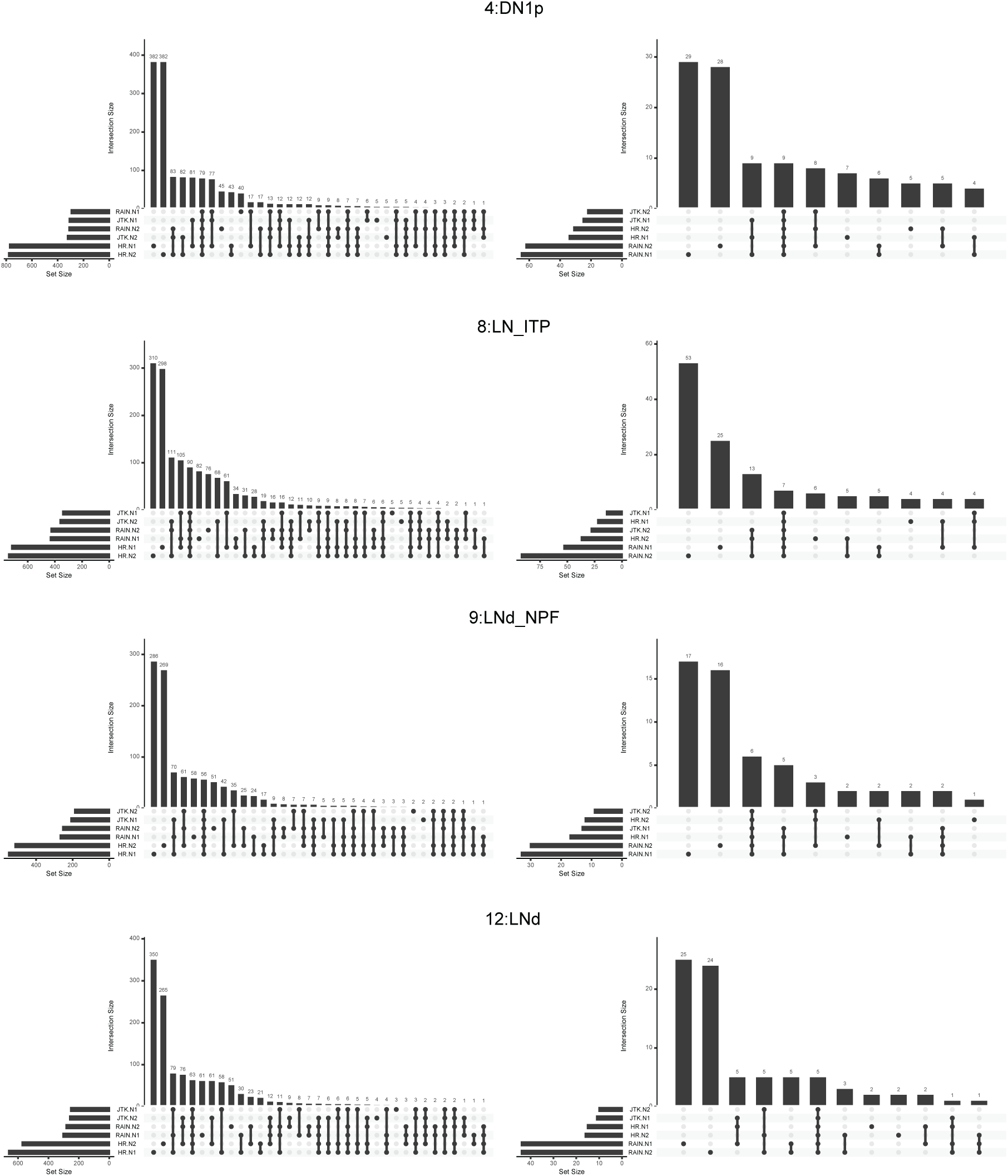
UpSet plots for cluster 4, 8, 9, and 12. The left (right) column uses *p*-values (adjusted *p*-values) to identify cycling genes.

**Figure S3:**
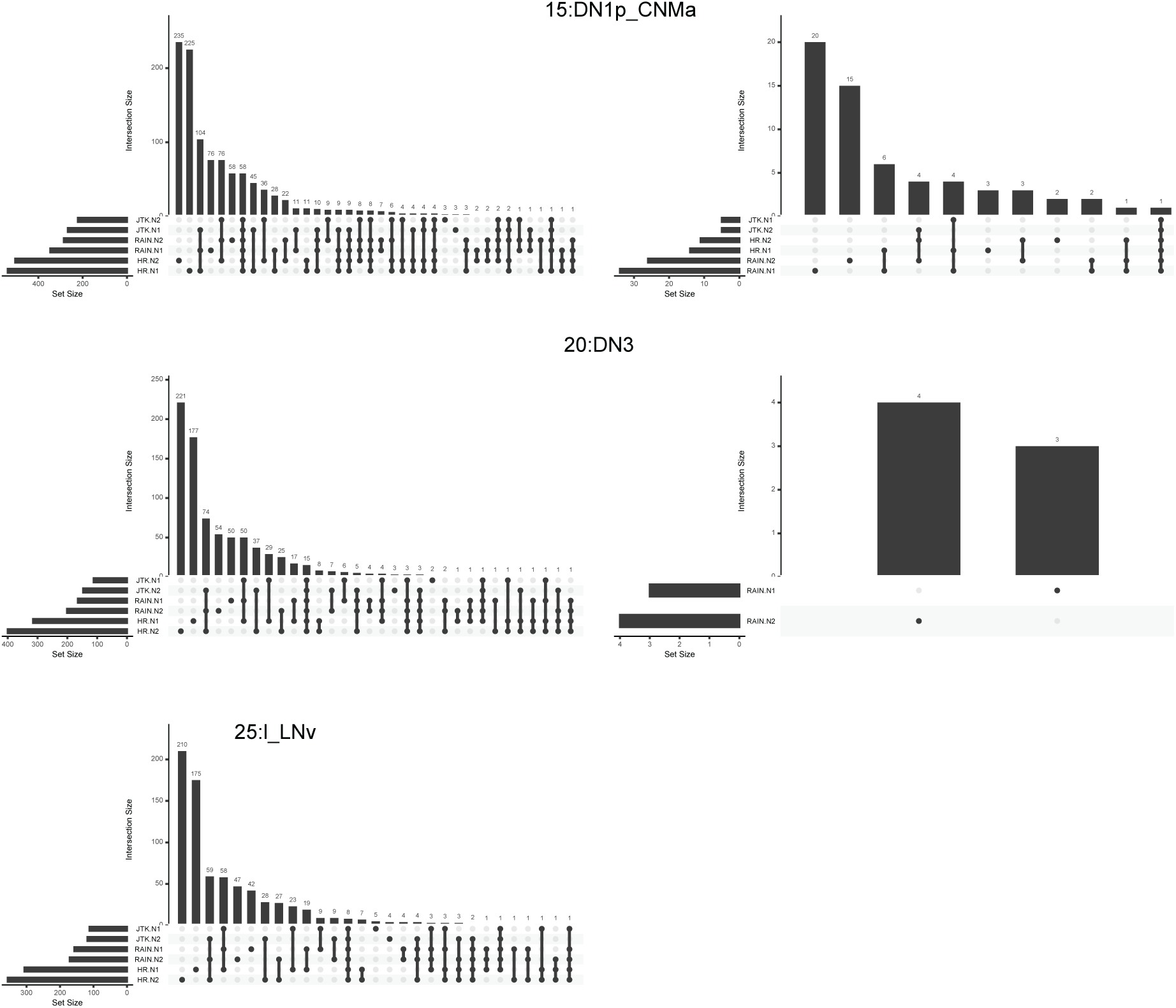
UpSet plots for cluster 15, 20, and 25. The left (right) column uses p-value (adjusted p-values) to identify cycling genes. Right column for cluster 25 is empty for that the average intersection between all sets were less than one.

**Figure S4:**
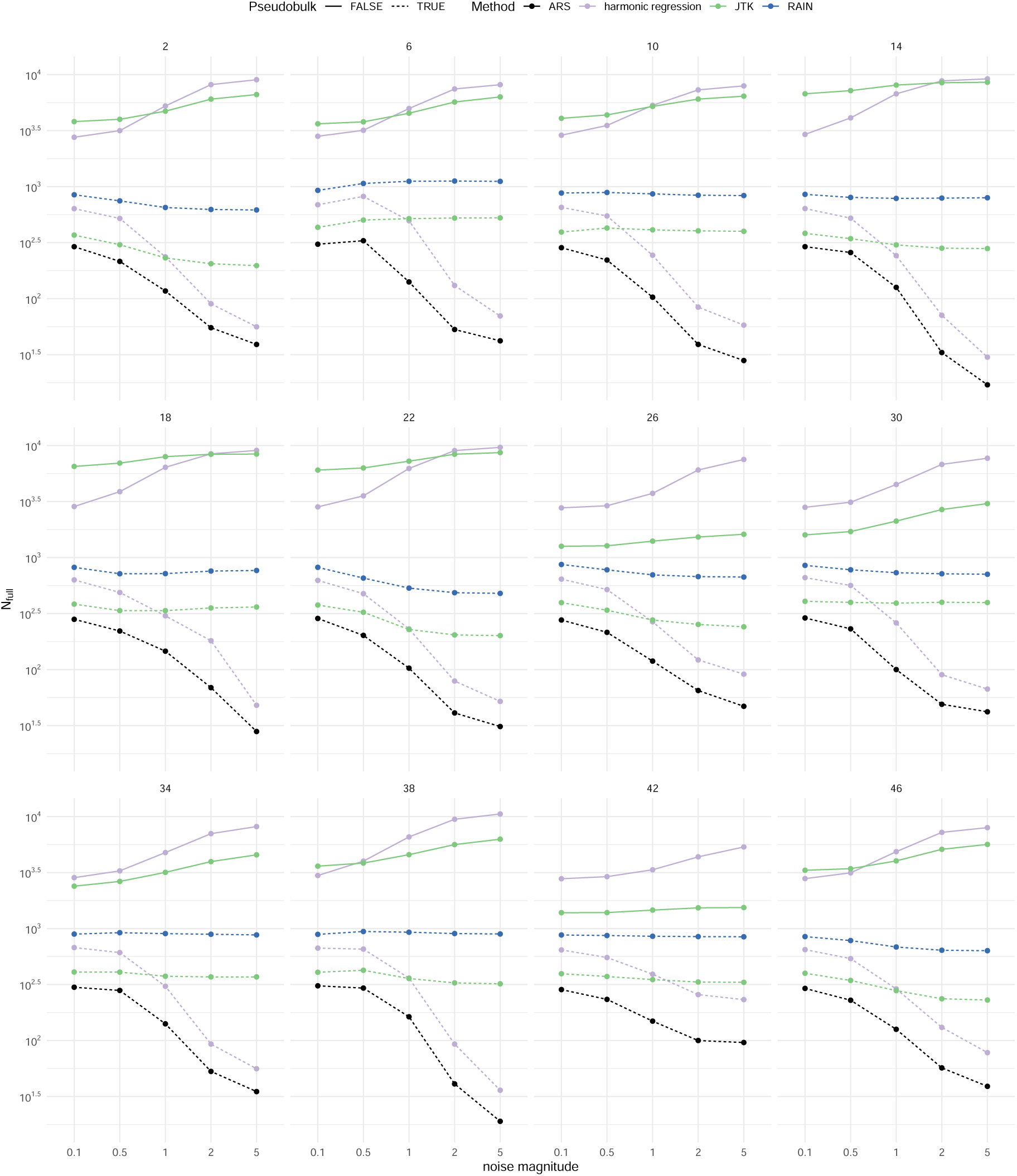
Effect of systematic contamination when using various methods with or without pseudo-bulking. Each panel shows how the number of detected cyclers *N*_full_ varies as function of the magnitude of noise injected at a single time-point.

**Figure S5:**
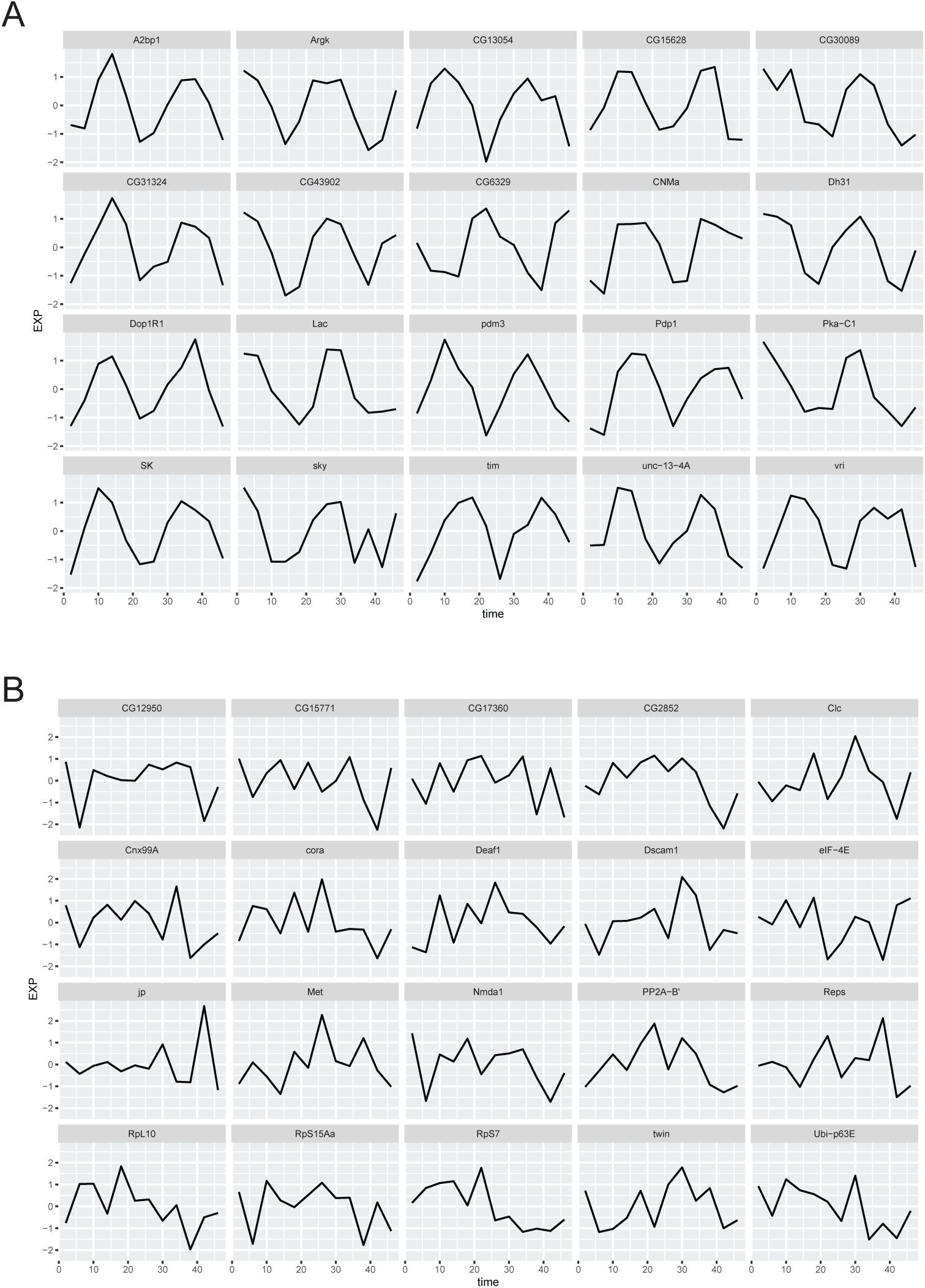
Rhythmic (A) and non-rhythmic (B) genes used for the generation of synthetic data.

**Figure S6:**
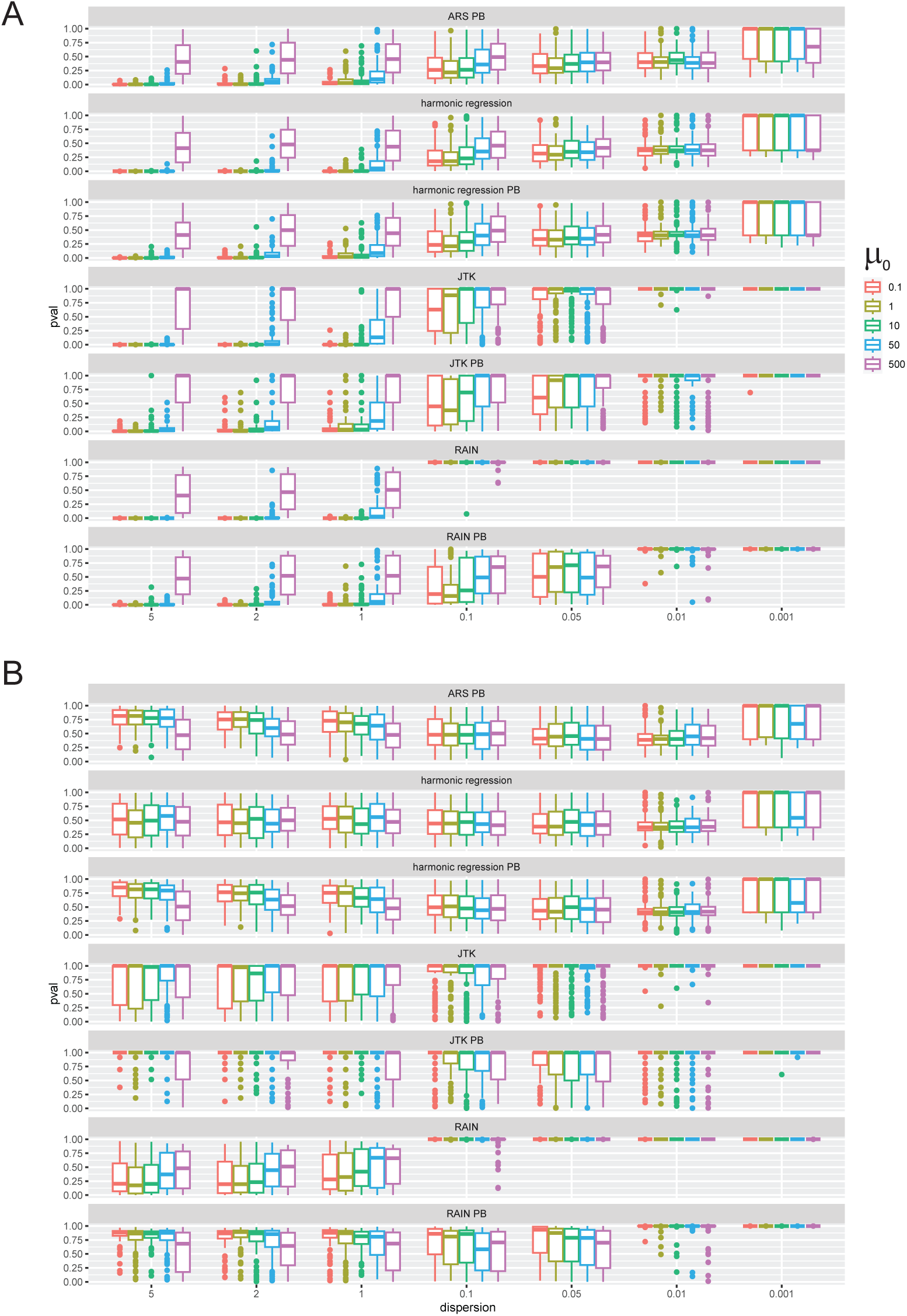
*p*-value distributions for cycling detection on synthetic rhythmic (A) and non-rhythmic (B) genes under different dispersions and µ0. Note that for (A) 0 is ideal and for (B) 1 is ideal. Dispersion goes from the largest to smallest so that variance increases from left to right.

**Figure S7:**
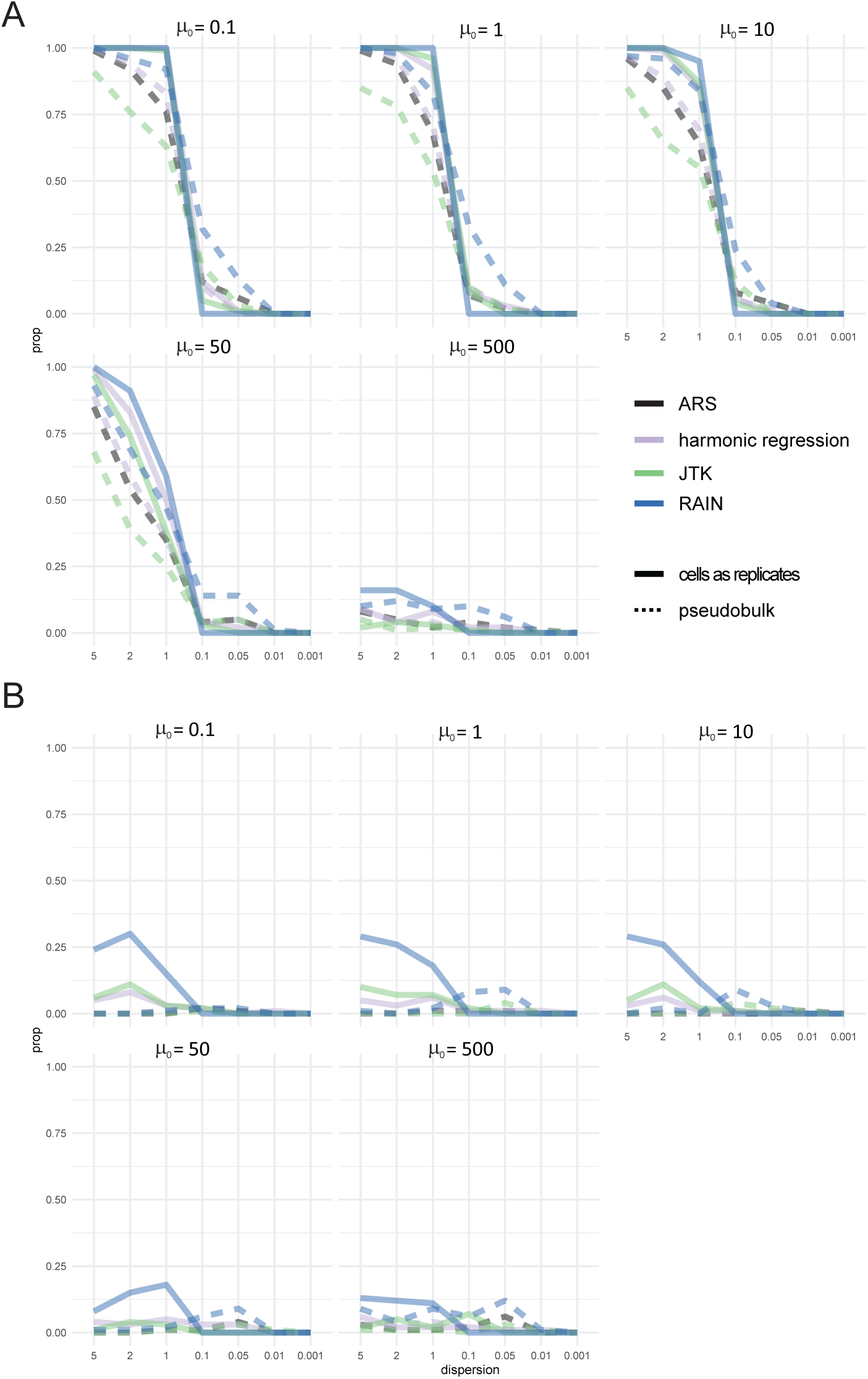
Cycling detection on synthetic rhythmic (A) and non-rhythmic (B) genes under different dispersion value and *µ*_0_. The *Y* -axis represents the overall proportion of synthetic genes that were considered to be cycling under each tested condition. The number at the top of each graph is the *µ*_0_ level. Note that for (A) 1 is ideal and for (B) 0 is ideal. Dispersion goes from the largest to smallest so that variance increases from left to right.

**Figure S8:**
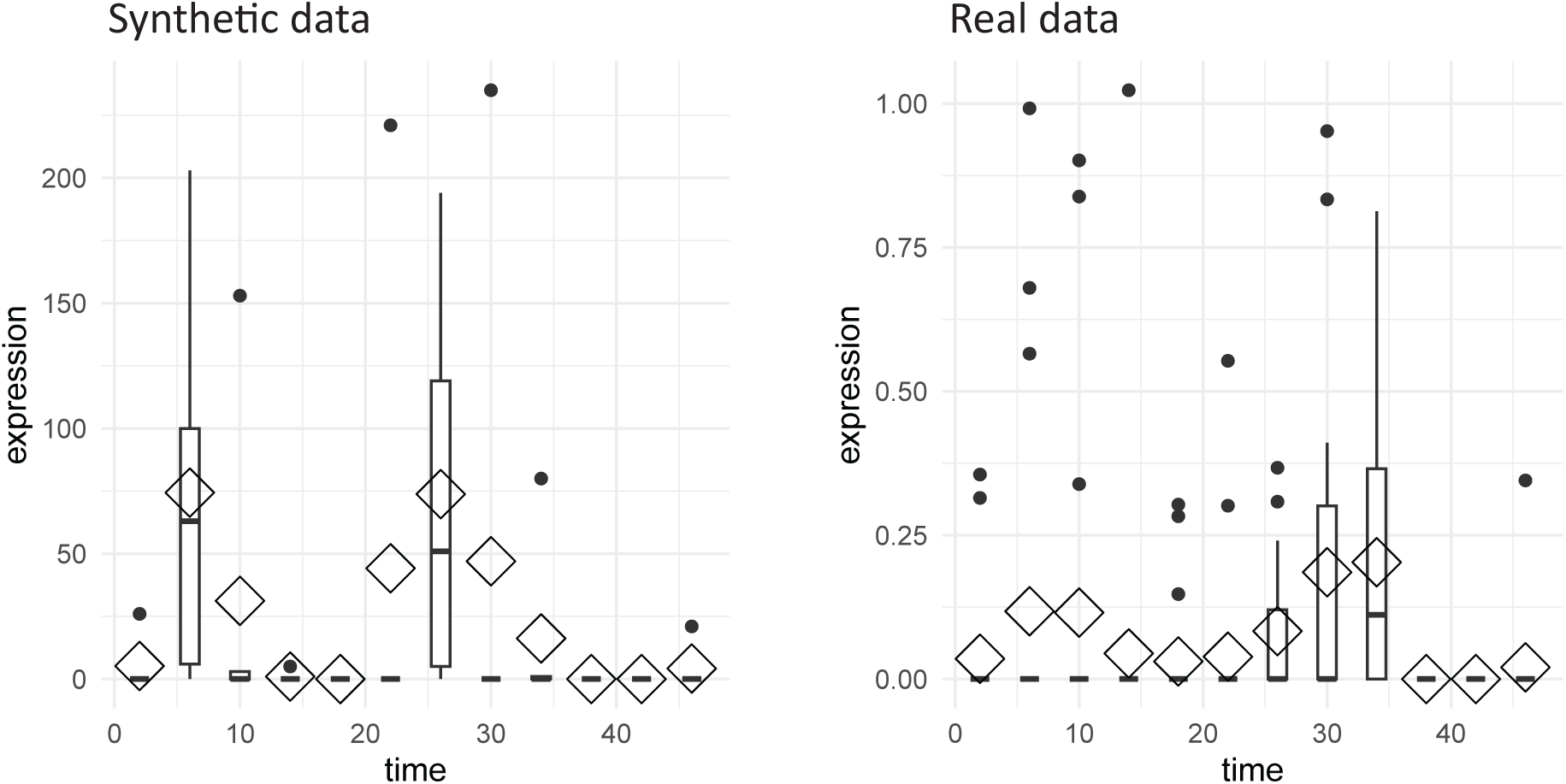
Example synthetic and real data (MsR1) data with oscillations in genes that are mostly unexpressed and are seemingly driven by noise/outliers. Here, diamonds represent means and heavy horizontal lines indicate medians.

**Figure S9:**
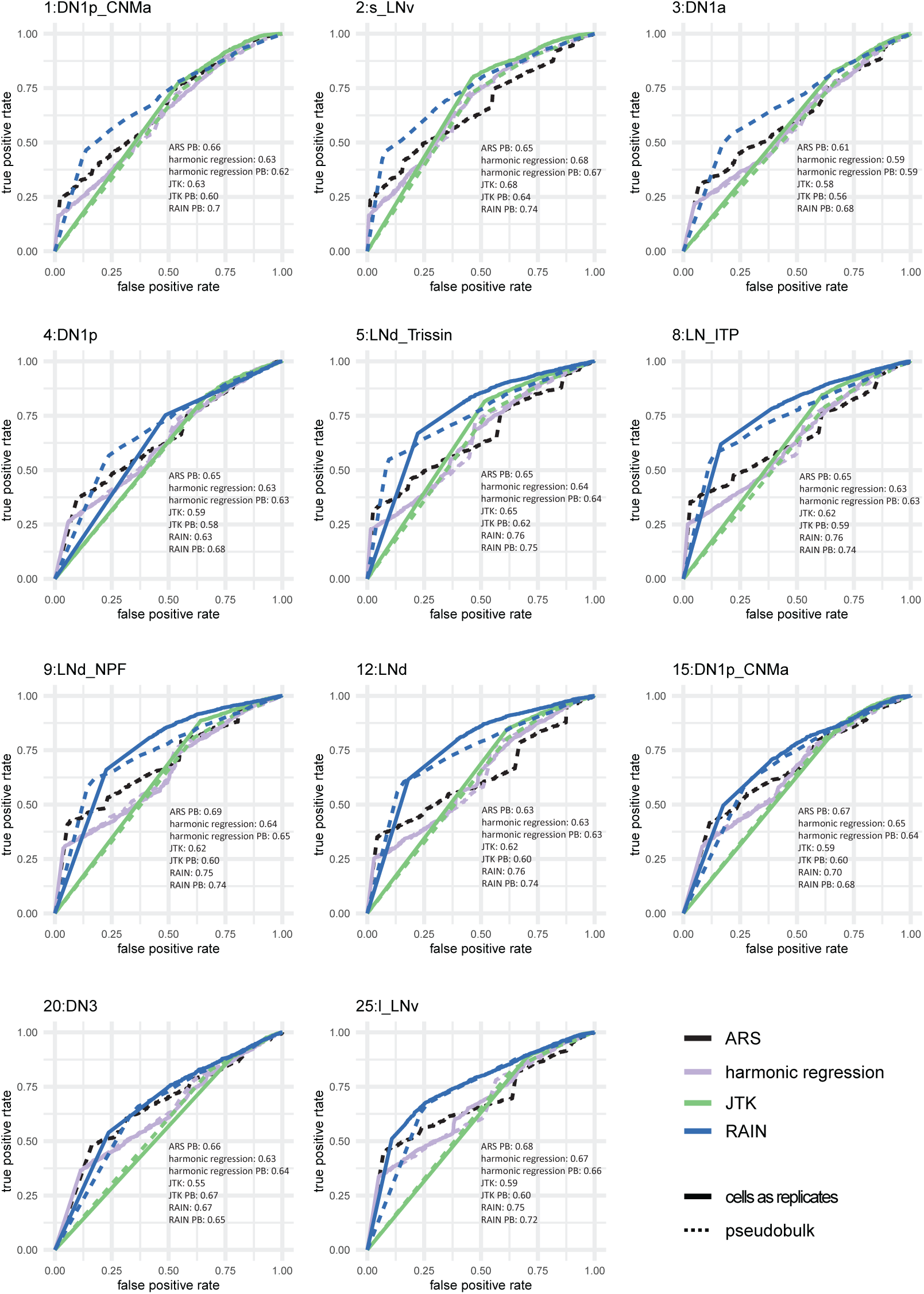
ROC curves of predicting circadian genes in bulk tissue using single cell RNA-seq data. All cells are supplied to the cycling detection algorithms and p-values are used for cycling prediction. l_LNv and s_LNv neurons were compared to bulk samples collected from the ventral lateral neurons. DN1p, DN1a neurons were compared to bulk samples collected from the dorsal neurons. LND neurons were compred to bulk samples collected from the dorsal lateral neurons. LN_ITP neurons were compared to samples taken from both the doral and ventral lateral neurons.

## Notes

### Competing Interest Statement

The authors have declared no competing interest.

### Summary of Updates

added details regarding comparing different ways to conduct cycling detection in synthetic data.

